# Immune mechanisms of type 1 diabetes revealed by single-cell transcriptomics, bulk transcriptomics, and experimental validation

**DOI:** 10.1101/2024.12.12.628291

**Authors:** Weisong Gao, Yue Zhu, Shuotong Zhang, Zhongming Wu

## Abstract

**Background:** Type 1 diabetes (T1D) is an autoimmune disorder characterized by the destruction of insulin-producing pancreatic β cells. Understanding the immune mechanisms underlying T1D is crucial for developing effective diagnostic and therapeutic strategies. This study aimed to elucidate the immune mechanisms of T1D by integrating single-cell RNA sequencing (scRNA-seq), bulk RNA-seq, and experimental validation.

**Methods:** scRNA-seq data (GSE200695) and bulk RNA-seq data (GSE9006) were obtained from the Gene Expression Omnibus (GEO) database. After data preprocessing, principal component analysis (PCA), and clustering, cell subtypes were annotated using ImmGenData as a reference. Receptor-ligand interactions were analyzed to identify key cell subtypes. Least absolute shrinkage and selection operator (LASSO) regression was performed to identify characteristic genes and construct a diagnostic model. Key genes were further validated using the training and validation sets. Functional enrichment and immune infiltration analyses were conducted for the key genes. In vitro experiments were performed to validate the findings using a high-glucose model in the monocytic cell line THP-1. siRNA-mediated knockdown of TRIB1 was performed to investigate its role in regulating monocyte activation and immune-related pathways under high-glucose conditions. Monocyte activation markers, inflammatory cytokines, apoptosis, and the expression of key genes and immune-related genes were assessed using immunofluorescence staining, ELISA, flow cytometry, qPCR, and Western blot.

**Results:** Monocytes were identified as the key cell subtype with the most interactions with other cell subtypes. Eleven characteristic genes were selected to construct a diagnostic model, which demonstrated high validation efficiency (AUC > 0.8). Three key genes (ACTG1, REL, and TRIB1) showed consistent expression trends in the training and validation sets. Functional analyses revealed that these genes were involved in immune regulation and PI3K/AKT/mTOR signaling. In vitro experiments confirmed that high glucose induced monocyte activation, as evidenced by increased expression of activation markers (CD86) and pro-inflammatory cytokines (IL-8 and TNF-α). High glucose also increased monocyte apoptosis and altered the expression of key genes (ACTG1, REL, and TRIB1) and immune-related genes (CXCL16, TGFBR1, CTLA4, CD48, TMIGD2, and HLA-DPB1). Knockdown of TRIB1 attenuated high glucose-induced monocyte activation, as demonstrated by decreased expression of activation markers and pro-inflammatory cytokines. TRIB1 knockdown also modulated the expression of immune-related genes and PI3K/AKT/mTOR signaling under high-glucose conditions.

**Conclusions:** This study integrates scRNA-seq, bulk RNA-seq, and experimental validation to unravel the immune mechanisms of T1D. Key genes (ACTG1, REL, and TRIB1) and monocytes were identified as crucial players in T1D pathogenesis. The constructed diagnostic model showed high validation efficiency. In vitro experiments confirmed the role of TRIB1 in regulating monocyte activation and immune-related pathways in a high-glucose model. These findings provide novel insights into the immune mechanisms of T1D and potential diagnostic and therapeutic targets.

## 1 Introduction

Type 1 diabetes (T1D) is a chronic autoimmune disorder characterized by the selective destruction of insulin-producing pancreatic β cells, resulting in hyperglycemia and severe complications ^[1]^. The incidence of T1D has been steadily increasing worldwide, particularly in children and adolescents, with an estimated 1.1 million individuals living with T1D in the United States alone ^[2]^. T1D is thought to result from a complex interplay between genetic susceptibility and environmental triggers, leading to the activation of autoreactive T cells and the production of autoantibodies against β cell antigens ^[3]^. The autoimmune attack on β cells leads to a progressive decline in insulin secretion, ultimately resulting in overt hyperglycemia and the need for exogenous insulin replacement therapy ^[4]^.

Despite advances in understanding the pathogenesis of T1D, the precise immune mechanisms underlying the disease remain poorly understood. T1D is characterized by a complex immune cell infiltration in the pancreatic islets, involving CD4+ and CD8+ T cells, B cells, macrophages, and dendritic cells ^[5–7]^. These immune cells interact with each other and with β cells through various cytokines, chemokines, and other signaling molecules, creating a proinflammatory microenvironment that promotes β cell damage and apoptosis ^[8, 9]^. However, the specific roles of different immune cell subtypes and their crosstalk in the pathogenesis of T1D have not been fully elucidated.

Recent advances in single-cell RNA sequencing (scRNA-seq) technology have revolutionized our understanding of cellular heterogeneity and cell-type-specific gene expression in complex tissues ^[10]^. By capturing the transcriptome of individual cells, scRNA-seq enables the identification of rare cell populations and the discovery of novel cell subtypes that may play crucial roles in disease pathogenesis ^[11]^. Integration of scRNA-seq data with bulk RNA-seq data provides a comprehensive view of the immune landscape and allows for the identification of key cell subtypes and genes involved in T1D ^[12]^.

Several studies have employed scRNA-seq to investigate the immune cell composition and gene expression profiles in T1D, focusing on pancreatic islets and the pancreatic lymph nodes ^[12–14]^. These studies have revealed the presence of distinct immune cell subtypes, such as CD8+ T cell subsets with different activation states and cytokine production profiles, and have identified novel gene signatures associated with T1D progression ^[14]^. However, the functional relevance of these findings and their potential as diagnostic or therapeutic targets have not been fully explored.

In the present study, we aimed to elucidate the immune mechanisms of T1D by integrating scRNA-seq data with bulk RNA-seq data and experimental validation. We first identified key cell subtypes and genes involved in T1D pathogenesis using bioinformatics analyses of publicly available datasets. We then constructed a diagnostic model based on the selected characteristic genes and evaluated its performance in both training and validation sets. Furthermore, we performed in vitro experiments using a high-glucose model in monocytes, a key cell subtype identified in our analyses, to investigate the effects of high glucose on monocyte activation, immune-related pathways, and the role of a key gene, TRIB1, in regulating these processes. Our findings provide novel insights into the immune mechanisms of T1D, contribute to a better understanding of the complex immune landscape, and highlight potential diagnostic biomarkers and therapeutic targets, paving the way for future research and clinical applications.

## 2 Materials and methods

### 2.1 Data sources and preprocessing

The single-cell RNA sequencing (scRNA-seq) data of GSE200695 and the bulk RNA-seq data of GSE9006 were obtained from the Gene Expression Omnibus (GEO) database. GSE200695 included 4 control samples and 6 T1D samples, while GSE9006 contained expression profile data and clinical information of 105 subjects (24 control individuals and 81 T1D patients) from both GPL96 and GPL97 platforms. T1D-related genes were retrieved from the GeneCards database.

The scRNA-seq expression file was analyzed using the Seurat package in R. Lowly expressed genes (nFeature_RNA > 500 and nFeature_RNA < 4000) were removed, and the data were subjected to dimensionality reduction using principal component analysis (PCA) and clustered using the t-distributed stochastic neighbor embedding (t-SNE) algorithm. Cell subtypes were annotated using ImmGenData as a reference and the R package "SingleR". Differentially expressed marker genes among various cell types were identified using the "FindAllMarkers" program with p_val_adj < 0.05 and |avg_log2FC| > 0.25 as thresholds.

### 2.2 Identification of key cell subtypes

Ligand-receptor interactions in the single-cell expression profile were statistically analyzed using the CellPhoneDB software package. The cluster labels of all cells were randomly permuted 10 times, and the mean expression levels of receptors in clusters and ligands in interacting clusters were determined. For each receptor-ligand pair in each pairwise comparison between two cell types, a null distribution was generated. Finally, ligand-receptor pairs of interest were selected for visualization, and monocytes were identified as the key cell subtype for further analysis.

### 2.3 Functional enrichment analysis and PPI network analysis

The Metascape database (www.metascape.org) was used for functional annotation of the marker genes identified in monocytes, the key cell subtype selected for further analysis. Gene Ontology (GO) and Kyoto Encyclopedia of Genes and Genomes (KEGG) pathway analyses were performed to explore the biological processes, molecular functions, cellular components, and signaling pathways associated with these marker genes. Results with a minimum overlap of 3 and p ≤ 0.01 were considered statistically significant. The protein-protein interaction network of these marker genes was constructed using the STRING database (https://string-db.org/) and visualized using Cytoscape software (version 3.8.2).

### 2.4 Diagnostic model construction

Characteristic genes were identified using the least absolute shrinkage and selection operator (LASSO) regression analysis with the glmnet package in R. A diagnostic model was constructed based on the selected characteristic genes, and its performance was evaluated using receiver operating characteristic (ROC) curve analysis in the training and validation sets. Key genes were further identified based on their consistent expression trends and significant differential expression between T1D and control samples in both the training and validation sets.

### 2.5 Immune infiltration

Immune cell infiltration in disease and control samples was assessed using the CIBERSORT algorithm, a computational method that estimates the relative abundance of different immune cell types within a mixed cell population based on gene expression data ^(^[15]^). Pearson’s correlation coefficients were calculated to determine the correlations among various immune cell types and visualized using a heatmap, enabling the identification of co-occurring or mutually exclusive immune cell populations. The Wilcoxon rank-sum test was employed to compare the proportions of immune cells between T1D and control samples, facilitating the identification of significantly altered immune cell populations in T1D. Furthermore, Spearman’s correlation analysis was performed to investigate the associations between the expression levels of key genes and various immune factors, including immunomodulators, chemokines, immunosuppressors, and major histocompatibility complex (MHC) molecules.

### 2.6 Gene set variation analysis (GSVA)

GSVA was performed to identify specific signaling pathways enriched by key genes using the GSVA package in R. The input data for GSVA included a normalized gene expression matrix and predefined gene sets from the Molecular Signatures Database (MSigDB). Differential pathway analysis was then performed using the limma package to determine significantly enriched signaling pathways in samples with high expression of key genes compared to those with low expression.

### 2.7 Regulatory network and ceRNA network analyses

Transcription factors were predicted using the R package "RcisTarget" based on motifs. The normalized enrichment score (NES) for each motif was calculated using the motif-gene set AUC and the distribution of AUCs across all motifs. The rcistarget.hg19.motifdb.cisbpont.500bp was used as the gene-motif rankings database. The competing endogenous RNA (ceRNA) network of key genes was constructed using the miRWalk, miRDB, and TargetScan databases and visualized using Cytoscape software.

### 2.8 In vitro experimental validation

#### 2.8.1 Monocyte cell culture and treatment

The human monocytic cell line THP-1 was obtained from the Cell Resource Center of the Shanghai Institute of Biological Sciences, Chinese Academy of Sciences. Cells were cultured in RPMI-1640 medium (Gibco, Grand Island, NY, USA) supplemented with 10% FBS (Gibco), 100 U/mL penicillin (Gibco), and 100 μg/mL streptomycin (Gibco) at 37℃ in a humidified atmosphere containing 5% CO_2_. To mimic hyperglycemic conditions, THP-1 cells were seeded at a density of 1 × 10^6^ cells/mL and treated with normal (5.5 mM) or high (30 mM) glucose for 24 or 48 h.

#### 2.8.2 siRNA transfection

THP-1 cells were transfected with small interfering RNA (siRNA) targeting TRIB1 or negative control siRNA (RiboBio, Guangzhou, China) using Lipofectamine 2000 (Invitrogen, Carlsbad, CA, USA) according to the manufacturer’s instructions. The siRNA sequences were as follows: si-TRIB1, 5’-GCAGCAGAAGAAACCCTTT-3’; si-NC, 5’-UUCUCCGAACGUGUCACGU-3’. The knockdown efficiency was confirmed by quantitative real-time PCR (qPCR) and Western blot. After transfection, THP-1 cells were cultured in normal glucose or high glucose conditions for 24 hours, and the effects of TRIB1 knockdown on THP-1 cell activation and immune-related pathways were evaluated.

#### 2.8.3 Immunofluorescence staining

THP-1 cells were fixed with 4% paraformaldehyde, permeabilized with 0.1% Triton X-100, and blocked with 1% bovine serum albumin (BSA). The cells were then incubated overnight at 4°C with primary antibodies against monocyte activation markers: CD86 (1:100, Abcam, Cambridge, UK). After washing, the cells were incubated with Alexa Fluor 488-conjugated secondary antibodies (1:500, Invitrogen) for 1 hour at room temperature. Nuclei were stained with DAPI (Invitrogen), and images were captured using an Olympus fluorescence microscope.

#### 2.8.4 Enzyme-linked immunosorbent assay (ELISA)

Levels of interleukin-8 (IL-8) and tumor necrosis factor-α (TNF-α) in the supernatants of THP-1 cell cultures were quantified using ELISA kits (R&D Systems, Minneapolis, MN, USA) ccording to the manufacturer’s guidelines. Briefly, cell-free supernatants and standards were added to antibody-coated wells and incubated for 2 h at room temperature. Following washing, detection antibodies were added and incubated for another 2 h. After additional washing, the substrate solution was added, and the reaction was stopped using the stop solution. Absorbance was read at 450 nm on a microplate reader (BioTek, Winooski, VT, USA).

#### 2.8.5 Flow cytometry

Apoptosis in THP-1 cells was evaluated using an Annexin V-FITC/PI kit (BD Biosciences, San Jose, CA, USA) per the manufacturer’s protocol. In brief, cells were collected, washed with cold PBS, and resuspended in binding buffer. Following staining with Annexin V-FITC and PI for 15 min at room temperature in the dark, the proportion of apoptotic cells was quantified by flow cytometry (BD FACSCalibur, BD Biosciences).

#### 2.8.6 Quantitative real-time PCR (qPCR)

Total RNA was extracted from THP-1 cells using TRIzol reagent (Invitrogen) and reverse-transcribed into cDNA using a PrimeScript RT Reagent Kit (TaKaRa, Kusatsu, Japan). qPCR was performed using SYBR Green Master Mix (TaKaRa) on a LightCycler 480 System (Roche, Basel, Switzerland). The primer sequences used for qPCR are listed in Supplementary Table 1. The expression levels of key genes (ACTG1, REL, and TRIB1), immune-related genes (CXCL16, TGFBR1, CTLA4, CD48, TMIGD2, and HLA-DPB1) were normalized to GAPDH and calculated using the 2^-ΔΔCt^ method.

#### 2.8.7 Western blot

THP-1 cells were lysed in RIPA buffer supplemented with protease and phosphatase (Beyotime, Shanghai, China). Protein samples were separated by SDS-PAGE, transferred to PVDF membranes (Millipore, Bedford, MA, USA), and blocked with 5% non-fat milk. The membranes were probed with primary antibodies specific for ACTG1 (1:1000, Abcam, Cambridge, UK), REL (1:1000, Cell Signaling Technology, Danvers, MA, USA), TRIB1 (1:1000, Abcam), PI3K (1:1000, Cell Signaling Technology, Danvers, MA, USA), phospho-PI3K (p-PI3K, 1:1000, Cell Signaling Technology), AKT (1:1000, Cell Signaling Technology), phospho-AKT (p-AKT, 1:1000, Cell Signaling Technology), mTOR (1:1000, Cell Signaling Technology), phospho-mTOR (p-mTOR, 1:1000, Cell Signaling Technology), and GAPDH (1:5000, Proteintech, Rosemont, IL, USA) at 4°C overnight. After incubation with HRP-conjugated secondary antibodies (1:5000, Cell Signaling Technology), the bands were visualized using an enhanced chemiluminescence system (Millipore).

### 2.9 Statistical analysis

Statistical differences between groups were determined using appropriate tests in GraphPad Prism 8.0. For two-group comparisons, an unpaired t-test was employed, while multiple groups were analyzed by one-way ANOVA with Tukey’s post hoc test. Results are presented as mean ± SD, and statistical significance was set at p < 0.05.

## 3 Results

### 3.1 Cluster analysis of cell subtypes identified with scRNA-seq

Quality control metrics, including the number of genes detected (nFeature_RNA), the number of unique molecular identifiers (nCount_RNA), and the percentage of mitochondrial genes (percent.mt), were evaluated for 83,348 cells with nFeature_RNA between 500 and 4000 (Fig. 1A). Principal component analysis (PCA) was conducted, and the relationship between samples based on the first two principal components was visualized (Fig. 1B). ElbowPlot determined the optimal number of principal components (PCs) to be 16 (Fig. 1C). t-distributed stochastic neighbor embedding (t-SNE) analysis identified 20 distinct cell subtypes (Fig. 1D). The 20 clusters were annotated into 6 cell categories using ImmGenData as the reference and the R package "SingleR": monocytes, innate lymphoid cells (ILCs), T cells, neutrophils, natural killer T (NKT) cells, and B cells (Fig. 1E). The FindAllMarkers program detected a total of 893 marker genes (Supplementary Table 1), which were used for further analysis.

**Figure 1.**
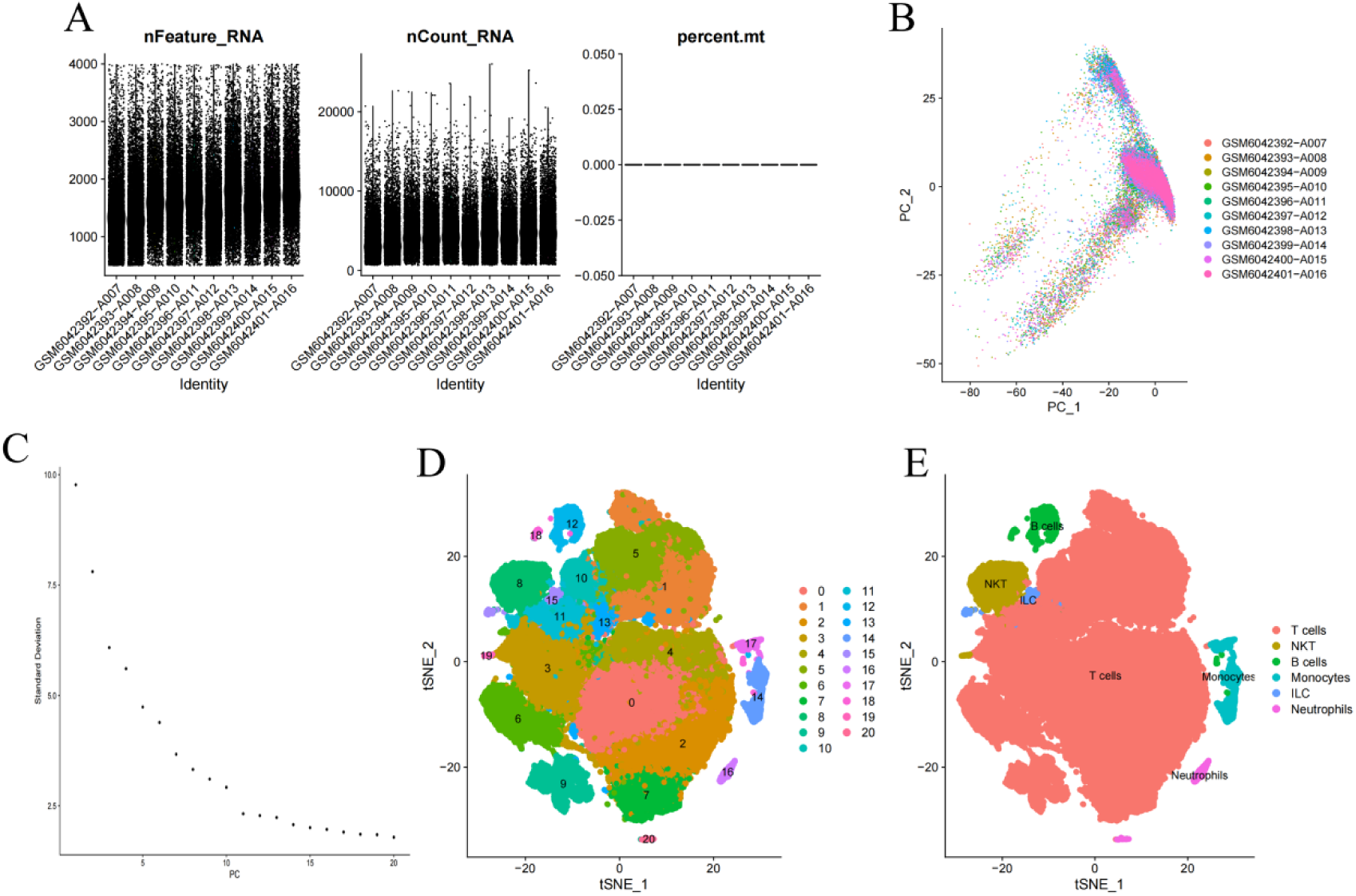
Quality control metrics and cell subtype identification using scRNA-seq data. (A) Quality control metrics for each cell in the single-cell RNA sequencing data. (B) Principal component analysis (PCA) plot showing the relationship between samples based on the first two principal components. (C) Elbow plot showing the optimal number of principal components (PCs). (D) t-distributed stochastic neighbor embedding (t-SNE) analysis identified 20 cell subtypes. (E) Annotation of cell subtypes using ImmGenData as the reference.

**Table 1.**
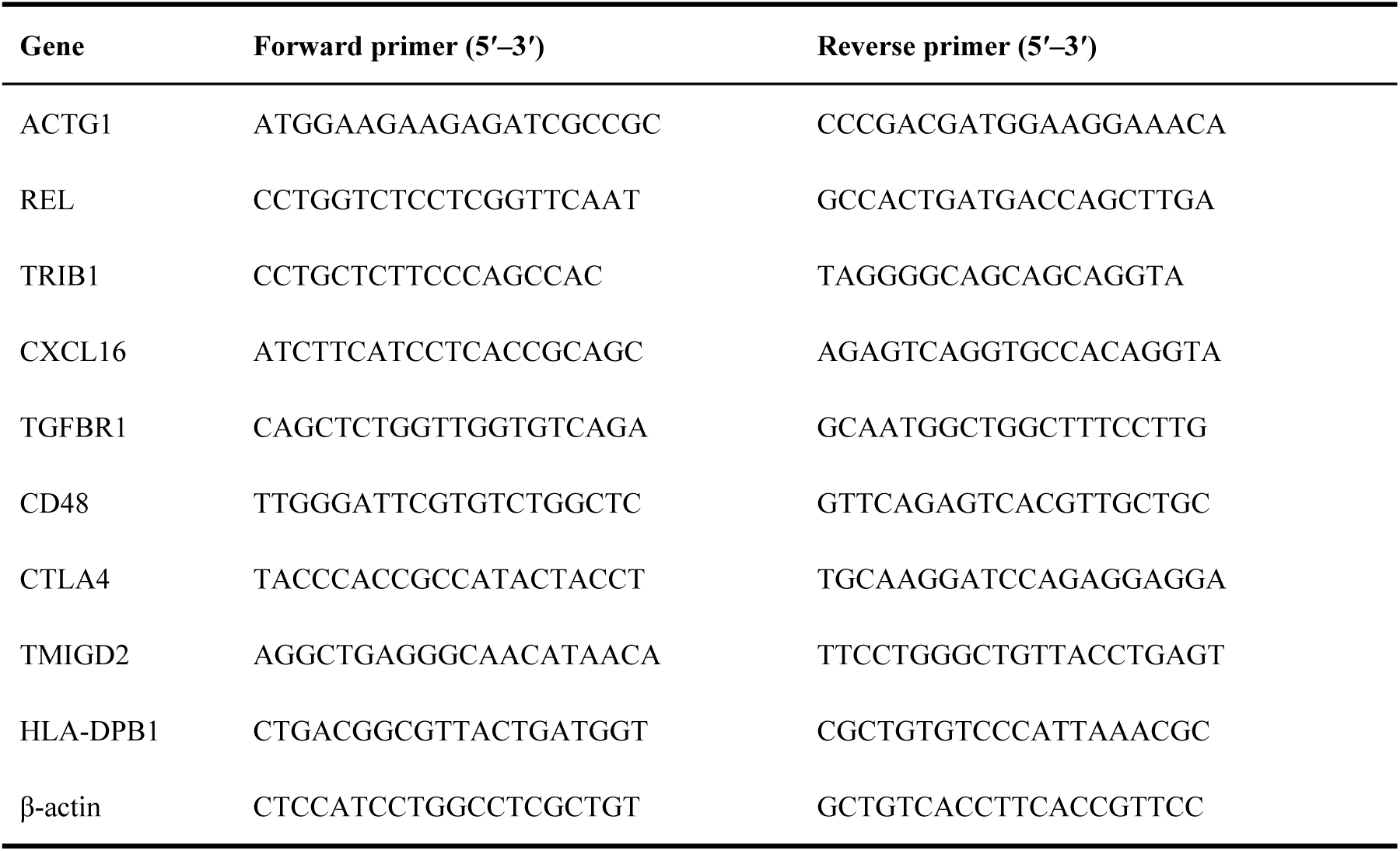
Primer sequences for q-PCR assay.

### 3.2 Analysis of receptor-ligand pairs

The analysis of ligand-receptor relationships suggested that monocytes|NKT and B cells|monocytes were significantly correlated with CD74_COPA and CD74_MIF (Fig. 2A). Moreover, more potential ligand-receptor pairs existed in neutrophils, monocytes, and ILCs (Fig. 2B). Receptor-ligand pairs were compared among these subtypes, revealing that the potential interaction of monocytes with other cell subtypes was the most complex (Fig. 2C). Accordingly, monocytes were selected, and their important role in T1D development was further analyzed.

**Figure 2.**
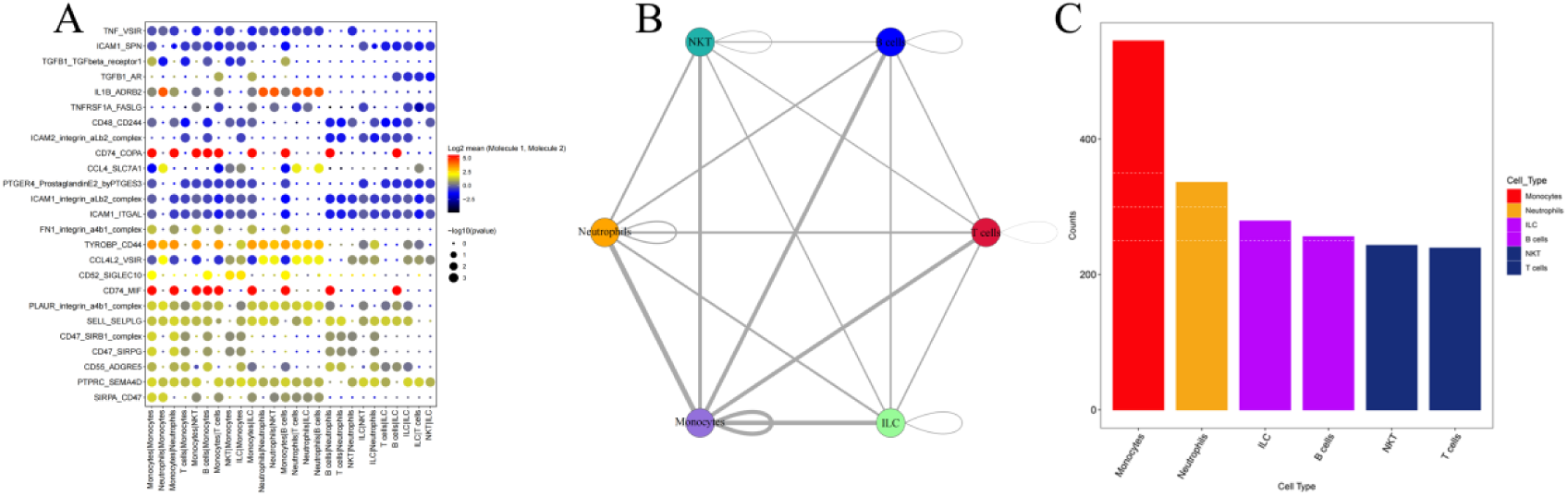
Receptor-ligand interaction analysis among cell subtypes. (A) Identification of significant ligand-receptor relationships between cell subtypes. (B) Quantification of potential ligand-receptor pairs in different cell subtypes. (C) Comparative analysis of receptor-ligand pairs among cell subtypes to determine the complexity of interactions.

### 3.3 Function analysis of marker genes for monocytes

A total of 315 marker genes for monocytes were screened. Functional enrichment analysis indicated that these marker genes were mainly enriched in several pathways, including negative regulation of immune system process, cell activation, regulation of cell activation, and PI3K-Akt signaling pathway (Fig. 3A). These marker genes were also used for protein-protein interaction network analysis using Cytoscape software (Fig. 3B).

**Figure 3.**
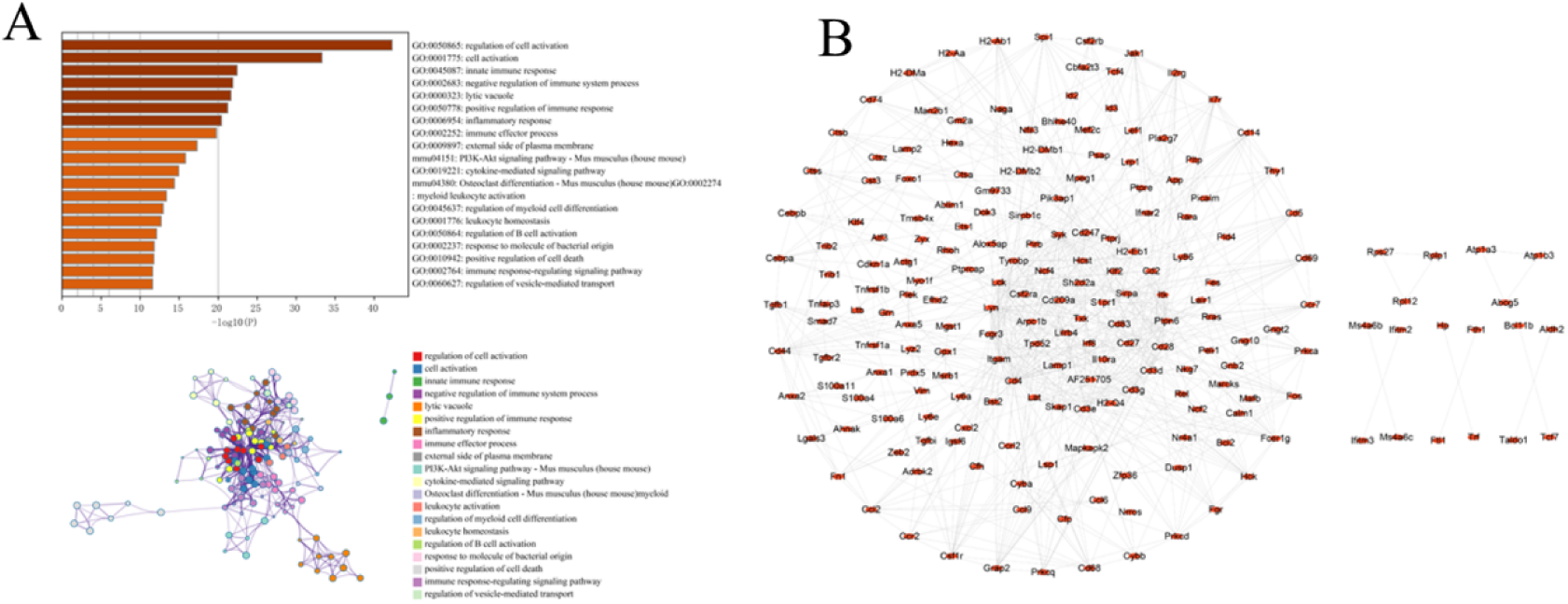
Functional characterization of marker genes in monocytes. (A) Functional enrichment analysis of marker genes in monocytes. (B) Protein-protein interaction network of marker genes in monocytes.

### 3.4 Construction of the diagnostic model and screening of characteristic genes

Subsequently, T1D-related characteristic genes were further identified by analyzing 315 marker genes of monocytes with the LASSO algorithm. A total of 11 characteristic genes were yielded, including PELI1, Actin Gamma 1 (ACTG1), RAP2A, REL, RHOH, CYBB, IL7R, MAFB, KLF2, TRIB1, and GIMAP6 (Fig. 4A). A risk-score model was derived according to the correlation coefficient of each gene (Fig. 4B). The stability and sensitivity of the risk model were confirmed with the receiver operating characteristic (ROC) curve, which unveiled that the area under the curve (AUC) values of the validation and training sets were 0.9465 and 0.8174, respectively, indicating the high validation efficiency of the risk model (Fig. 4C, 4D).

**Figure 4.**
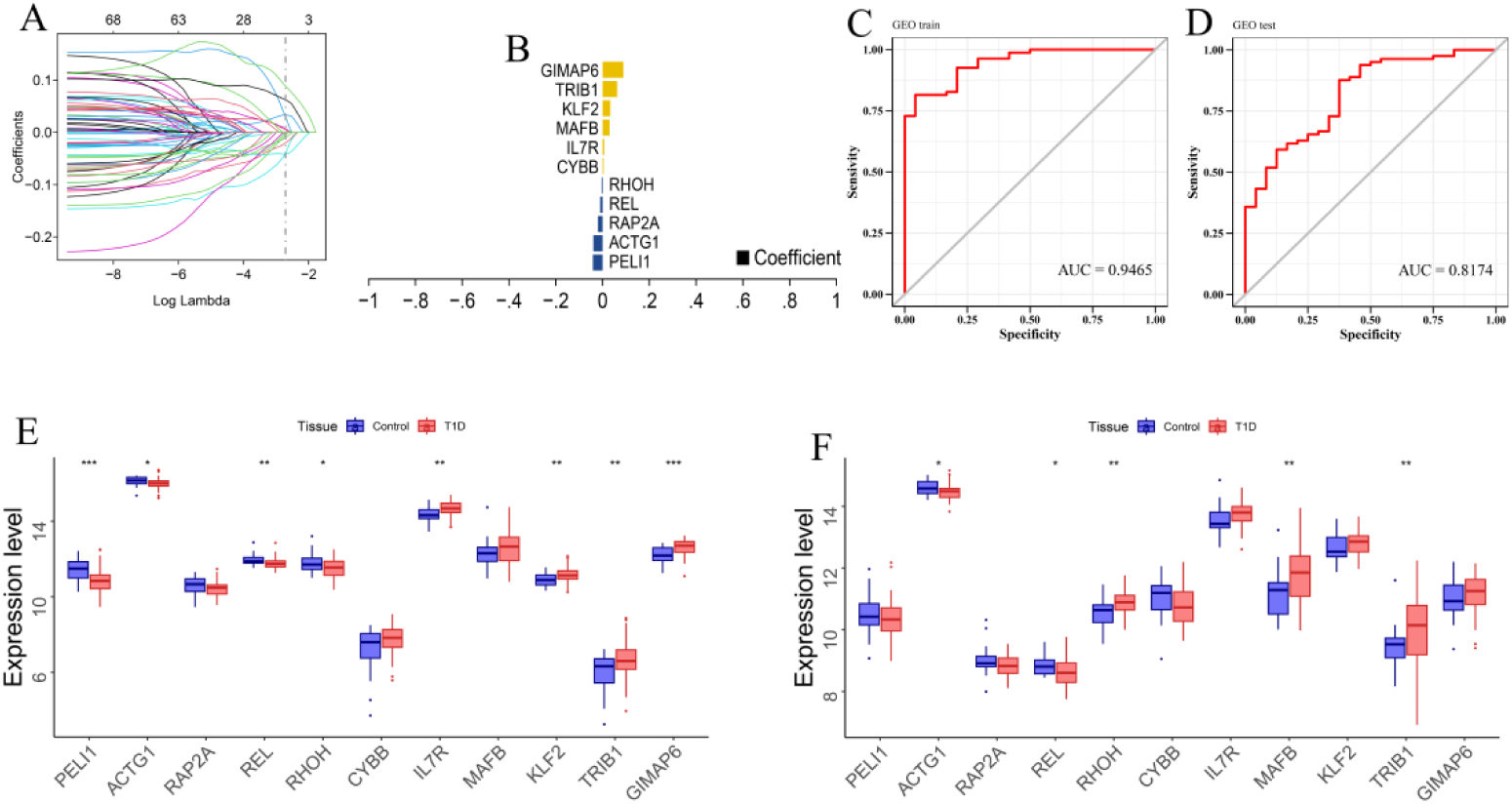
Diagnostic model construction and characteristic gene screening. (A) Identification of characteristic genes using the LASSO algorithm. (B) Development of a risk-score model based on the correlation coefficients of characteristic genes. (C, D) Evaluation of the stability and sensitivity of the risk model using receiver operating characteristic (ROC) curves in the (C) validation and (D) training sets. (E, F) Assessment of the expression patterns of characteristic genes in the (E) training and (F) validation sets.

To further screen for key genes, we examined the expression of the 11 characteristic genes. The results showed that only three characteristic genes (ACTG1, REL, and TRIB1) exhibited consistent expression trends and significant differential expression between T1D and control samples in both the training and validation sets (Fig. 4E, 4F). Consequently, these three characteristic genes (ACTG1, REL, and TRIB1) were considered potential key genes in T1D pathogenesis in our study. Furthermore, we analyzed the expression of these three key genes in each cell type using the single-cell dataset. The analysis revealed that TRIB1, REL and ACTG1 were all highly expressed in monocytes and neutrophils (Fig. 5).

**Figure 5.**
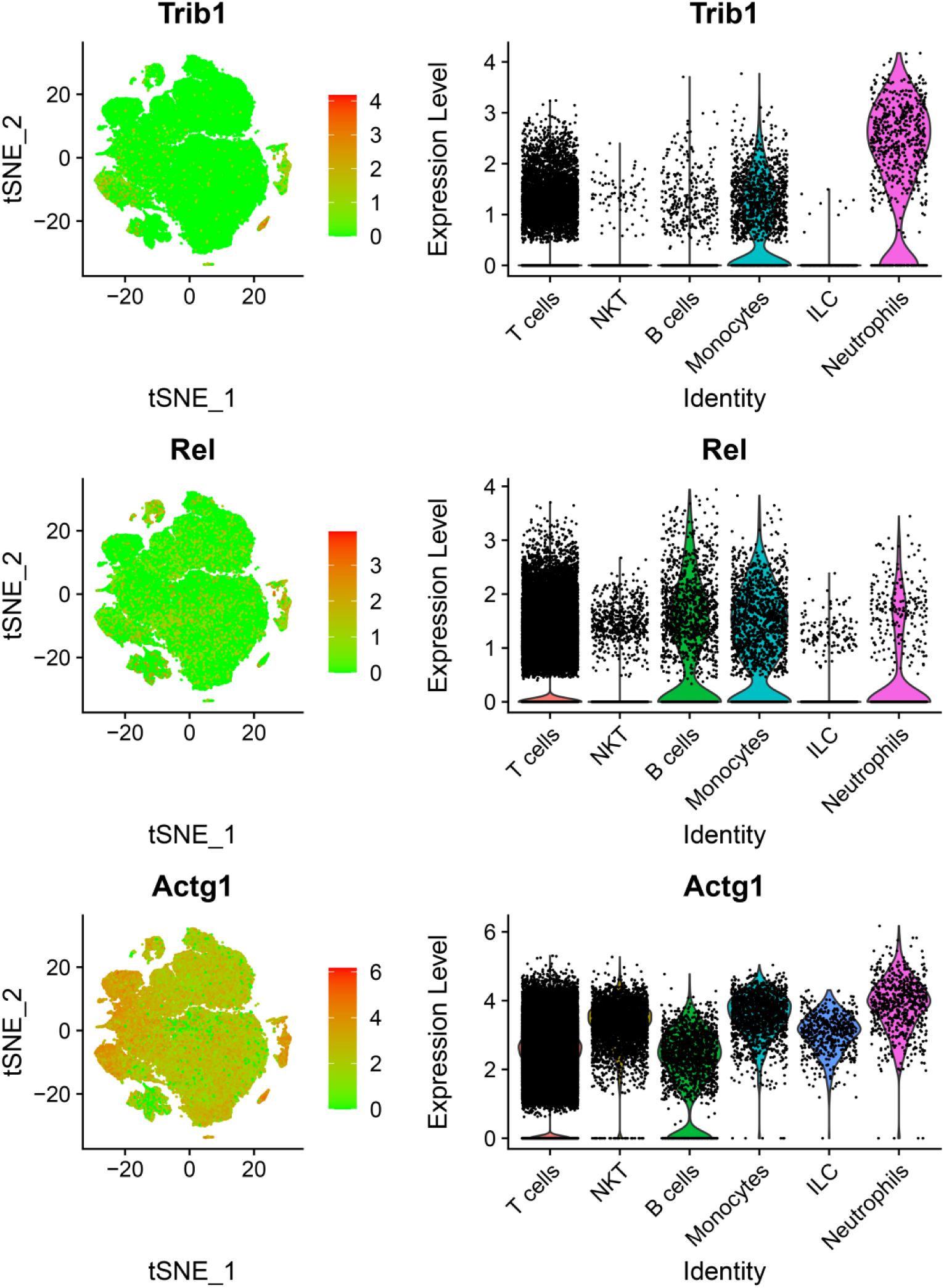
Cell type-specific expression of key genes. Visualization of the expression levels of TRIB1, REL, and ACTG1 across different cell types using the single-cell dataset.

### 3.5 Immune infiltration analysis

The relative proportions and correlations of each immune cell in disease and control samples are shown in Fig. 6A and 6B. In T1D samples, the level of resting memory CD4+ T cells and CD8+ T cells were significantly higher than that in normal patients, while the level of γδ T cells was significantly lower (Fig. 6C). Additionally, TRIB1 was negatively correlated with CD8+ T cells, and ACTG1 was positively correlated with neutrophils and negatively correlated with Macrophages M2, REL was positively correlated with activated Dendritic cells and negatively correlated with CD8+ T cells, indicating that the three key genes (ACTG1, REL, and TRIB1) were highly associated with immune cells (Fig. 6D). Further analysis of the correlations between ACTG1, REL, and TRIB1 and different immune factors (including chemokines, immunoinhibitor, immunostimulator, and MHC) showed that the key genes were significantly correlated with immune factors. Specifically, TRIB1 was significantly correlated with cytokines CXCL16, TGFBR1, CTLA4, CD48, TMIGD2, and HLA-DPB1; ACTG1 was significantly correlated with cytokines CXCL17, CXCL16, HAVCR2, IL6R, TNFSF15, TMEM173, CD48, and C10orf54; andREL was significantly correlated with cytokines CXCL16, BTLA, NT5E, PVR, TNFRSF18, TNFSF13B, COSLG, and TAP2 (Fig. 6E-6H). These analyses confirmed that the T1D-related key genes were closely associated with immune cell infiltration levels and played important roles in the immune microenvironment.

**Figure 6.**
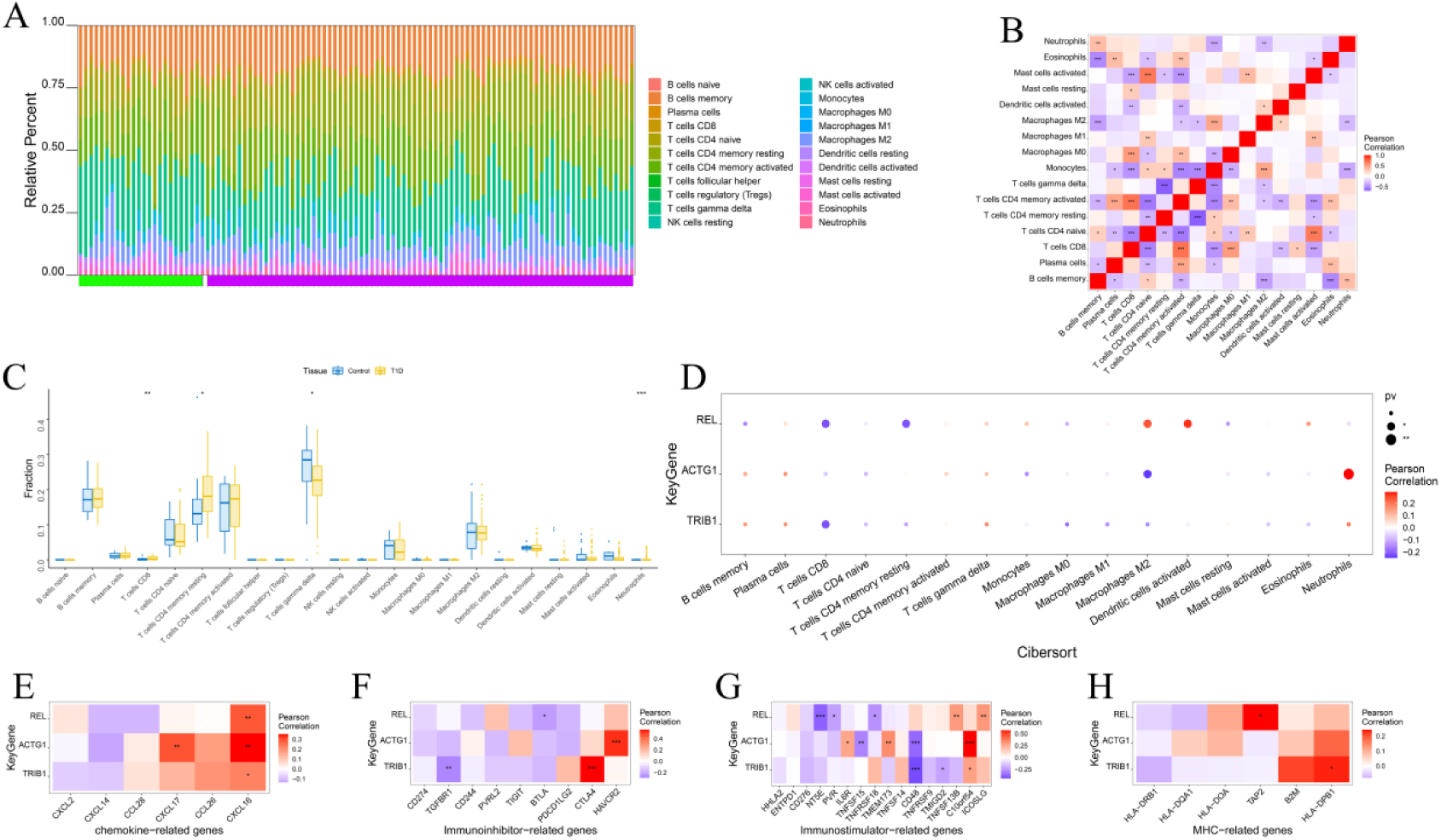
Analysis of immune cell infiltration and its relationship with key genes. (A) Quantification analysis of immune cell proportions in disease and control samples. (B) Correlation analysis of immune cell proportions in disease and control samples. (C) Comparative analysis of immune cell levels between T1D and control samples. (D) Assessment of the correlations between key genes (ACTG1, REL, and TRIB1) and immune cell proportions. (E-H) Investigation of the correlations between key genes and different immune factors, including (E)chemokines, (F) immunoinhibitor, (G) immunostimulator, and (H) MHC molecules.

### 3.6 Analysis of specific pathways related to key genes

Next, we employed gene set variation analysis (GSVA) to investigate the specific signaling pathways enriched by key genes. The results showed that TRIB1 expression was significantly positively correlated with 16 signaling pathways, including APOPTOSIS, TNFA SIGNALING VIA NFKB, and PI3K AKT MTOR SIGNALING, and negatively correlated with 2 signaling pathways, including HEME METABOLISM, ESTROGEN RESPONSE EARLY, and ESTROGEN RESPONSE LATE (Fig. 7A). ACTG1 expression was significantly positively correlated with 20 signaling pathways, including G2M CHECKPOINT, DNA REPAIR, and MITOTIC SPINDLE, and negatively correlated with 10 signaling pathways, including PI3K AKT MTOR SIGNALING, TNFA SIGNALING VIA NFKB, and PEROXISOME (Fig. 7B). REL expression was significantly positively correlated with 17 signaling pathways, including PROTEIN SECRETION, ADIPOGENESIS, and ANGIOGENESIS, and negatively correlated with 5 signaling pathways, including MYC TARGETS V2, TNFA SIGNALING VIA NFKB, and PI3K AKT MTOR SIGNALING (Fig. 7C).

**Figure 7.**
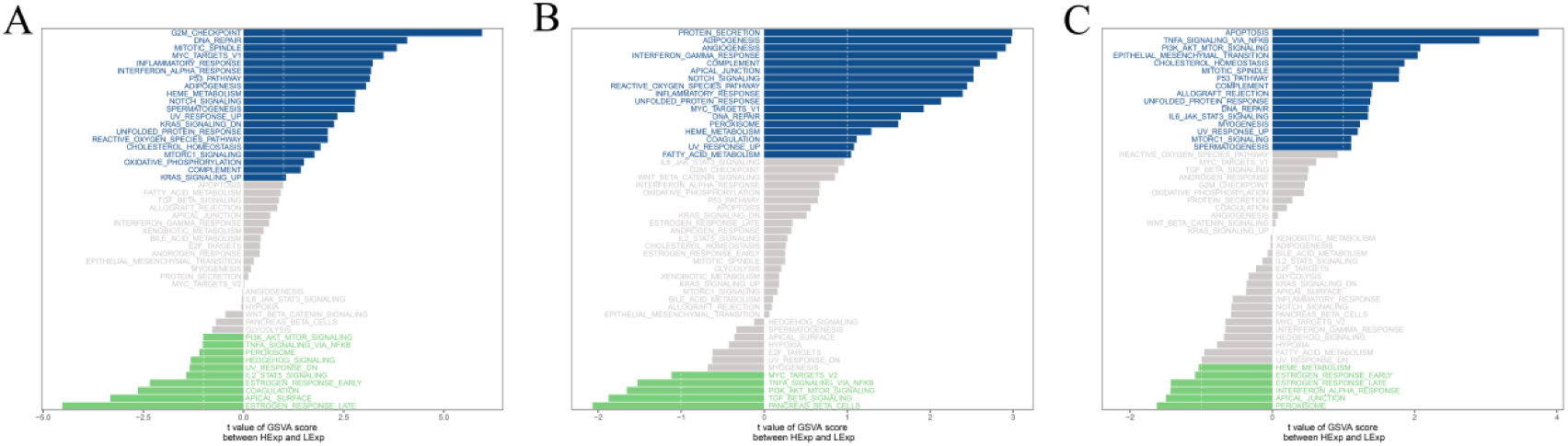
Analysis of specific pathways related to key genes. (A-C) Gene set variation analysis (GSVA) showing signaling pathways correlated with (A) ACTG1, (B) REL and (C) TRIB1 expression.

### 3.7 Regulatory network and ceRNA network analyses of key genes

Three key genes (ACTG1, REL, and TRIB1) were enriched in the motif cisbp M0367 (NES: 4.71) (Fig. 8A). A total of 249 mRNA-miRNA pairs were selected, and a network map was constructed using Cytoscape software. Compared with TRIB1 and ACTG1, more targeting miRNAs were associated with REL. Interestingly, there were four common targeting miRNAs between REL and TRIB1, including hsa-miR-4458, hsa-miR-4441, hsa-miR-574-5p, and hsa-let-7e-5p (Fig. 8B). This cross-over relationship warrants further study.

**Figure 8.**
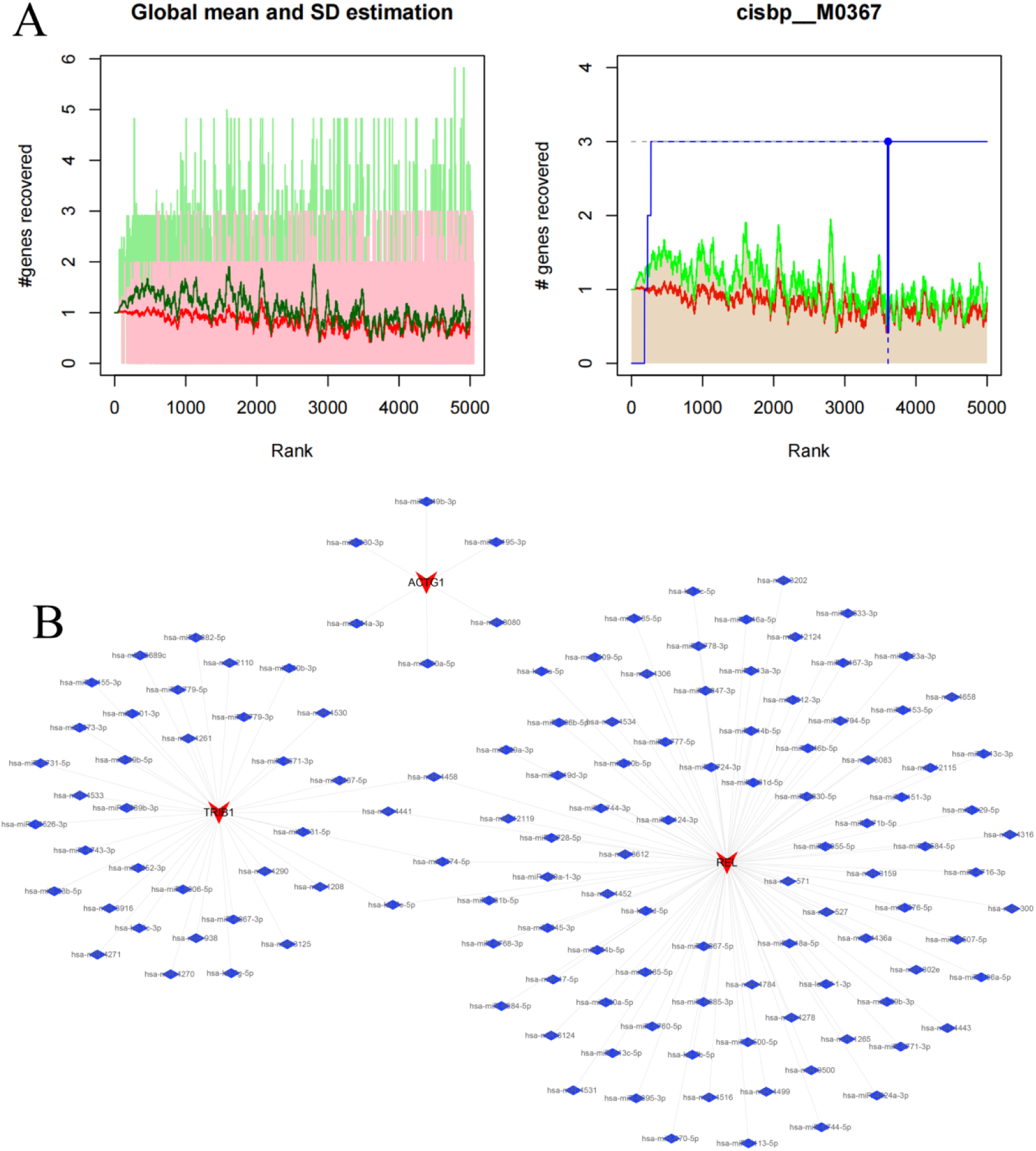
Regulatory and ceRNA network analyses of key genes. (A) Motif enrichment analysis of key genes to identify potential regulatory elements. (B) Construction of an mRNA-miRNA interaction network to visualize the targeting relationships between key genes and miRNAs.

### 3.8 High glucose induces monocyte activation and alters the expression of key genes

Furthermore, we employed a high-glucose treated monocyte THP-1 model to simulate a diabetes in vitro model. Immunofluorescence staining showed that high glucose (30 mM) significantly induced the expression of the THP-1 cell activation marker CD86 compared to normal glucose (5.5 mM) (Fig. 9A). ELISA results showed that high glucose significantly increased the secretion of proinflammatory cytokines, such as IL-8 and TNF-α, by THP-1 cells (p < 0.01) (Fig. 9B). Flow cytometry analysis indicated that high glucose significantly promoted THP-1 apoptosis, as evidenced by the increased percentage of Annexin V-positive cells (p < 0.001) (Fig. 9C). These data suggest that high glucose induces monocyte activation, which may contribute to the pathogenesis of T1D.

**Figure 9.**
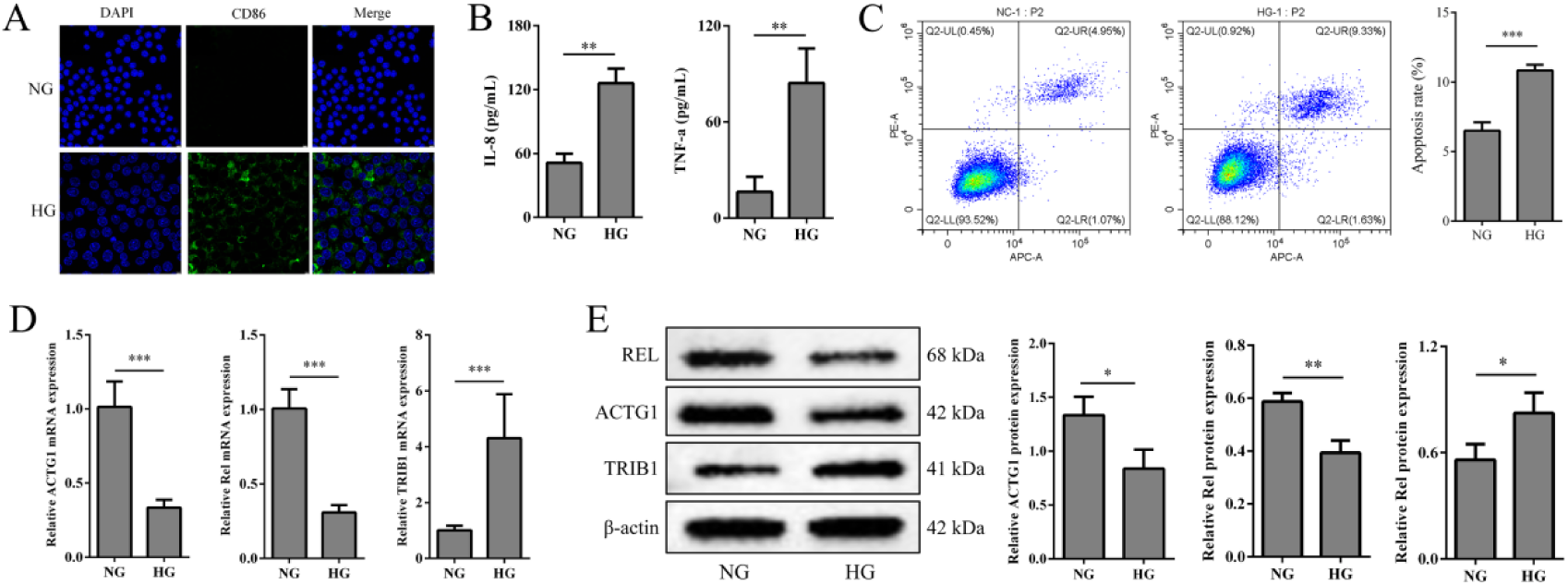
High glucose induces monocyte activation and alters the expression of key genes in THP-1 cells. (A) Immunofluorescence staining showing the expression of the monocyte activation marker CD86 in THP-1 cells treated with normal glucose (NG group) and high glucose (HG group). Scale bar: 50 μm. (B) ELISA analysis of proinflammatory cytokines IL-8 and TNF-α secretion by THP-1 cells treated with normal glucose and high glucose. (C) Flow cytometry analysis of THP-1 cell apoptosis using Annexin V staining in normal glucose and high glucose conditions. (D) qPCR analysis of the expression of key genes ACTG1, REL, and TRIB1 in THP-1 cells treated with normal glucose and high glucose. (E) Western blot analysis of the expression of key genes ACTG1, REL, and TRIB1 in THP-1 cells treated with normal glucose and high glucose. β-actin was used as a loading control. Data are presented as mean ± SD. Data are presented as mean ± SD. *p < 0.05, **p < 0.01.

Moreover, we performed expression validation of key genes by qPCR and Western blot. The results showed that high glucose significantly upregulated the expression of key genes ACTG1 and REL while downregulating TRIB1 expression in THP-1 cells (p < 0.05) (Fig. 9D, 10E). These findings indicate that high glucose modulates the expression of key genes, which may lead to immune dysregulation in T1D.

### 3.9 TRIB1 knockdown attenuates high glucose-induced monocyte activation and regulates immune-related pathways

To investigate the role of TRIB1 in regulating monocyte activation and immune-related pathways under high-glucose conditions, we performed siRNA-mediated TRIB1 knockdown in THP-1 cells. TRIB1 knockdown significantly attenuated high glucose-induced monocyte activation, as evidenced by the reduced expression of activation markers (CD86) (Fig. 10A) and proinflammatory cytokines (IL-8 and TNF-α) (p < 0.01) (Fig. 10B). Moreover, flow cytometry analysis revealed that high glucose treatment significantly increased the percentage of apoptotic THP-1 cells compared to the normal glucose group (p < 0.01). Notably, TRIB1 knockdown partially reversed the high glucose-induced apoptosis in THP-1 cells (p < 0.05) (Fig. 10C), suggesting that TRIB1 plays a role in regulating monocyte apoptosis under hyperglycemic conditions.

**Figure 10.**
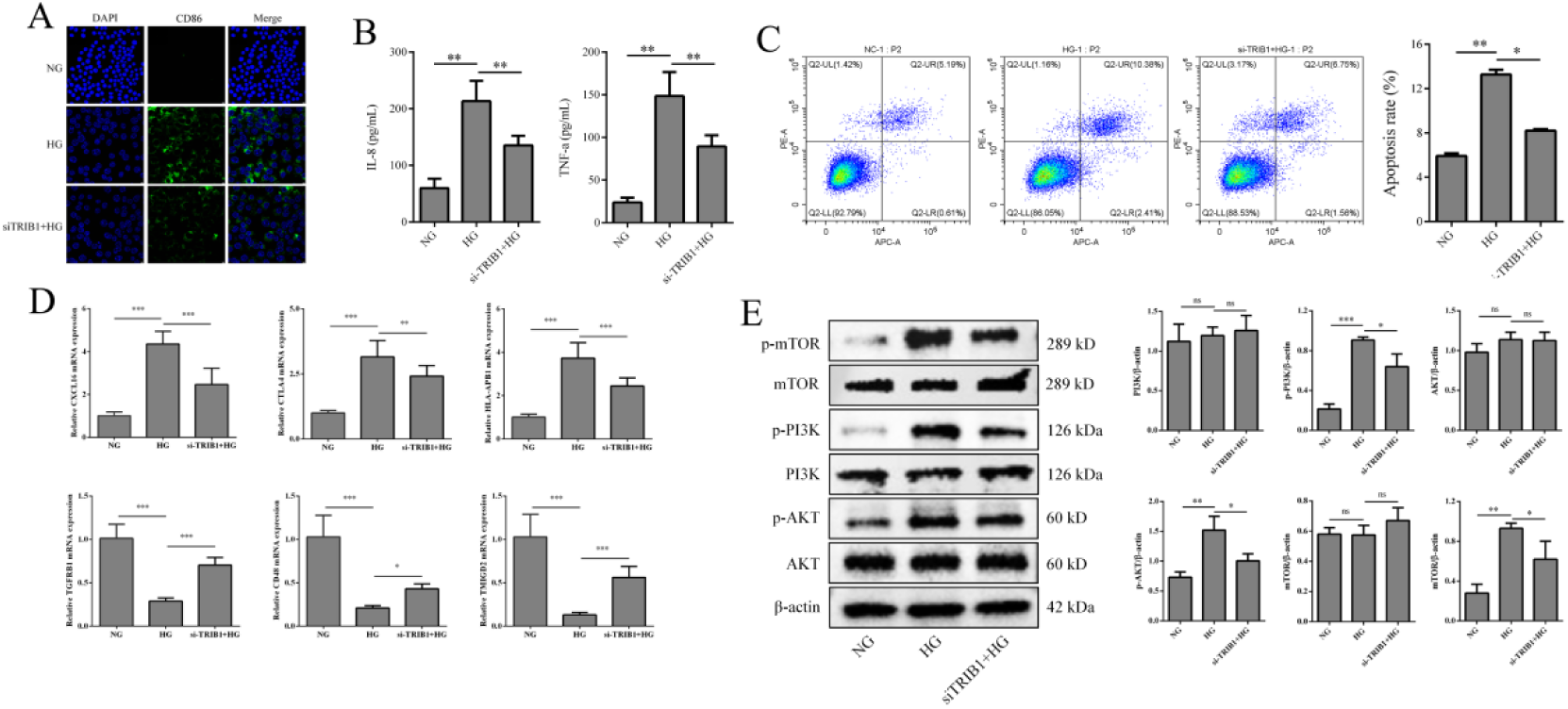
TRIB1 knockdown attenuates high glucose-induced monocyte activation and regulates immune-related pathways in THP-1 cells. (A) Immunofluorescence staining of the monocyte activation marker CD86 in THP-1 cells treated with normal glucose (NG), high glucose (HG), and high glucose with TRIB1 siRNA (si-TRIB1+HG). Scale bar: 50 μm. (B) ELISA analysis of IL-8 and TNF-α secretion by THP-1 cells in the NG, HG, and si-TRIB1+HG groups. (C) Flow cytometry analysis of THP-1 cell apoptosis using Annexin V staining in the NG, HG, and si-TRIB1+HG groups. (D) qPCR analysis of the mRNA expression levels of immune-related genes CXCL16, CTLA4, HLA-DPB1,TGFBR1, CD48 and TMIGD2 in THP-1 cells treated with NG, HG, and si-TRIB1+HG. (E) Western blot analysis of the protein expression levels of PI3K, p-PI3K, AKT, p-AKT, mTOR, and p-mTOR in THP-1 cells treated with NG, HG, and si-TRIB1+HG. β-actin was used as a loading control. Data are presented as mean ± SD. *p < 0.05, **p < 0.01. ns, not significant.

In the "Immune infiltration analysis" section, TRIB1 was significantly correlated with immune-related genes (CXCL16, TGFBR1, CTLA4, CD48, TMIGD2, and HLA-DPB1), suggesting that TRIB1 may regulate the expression of these immune-related genes. Therefore, we explored the effect of TRIB1 knockdown on the expression of CXCL16, TGFBR1, CTLA4, CD48, TMIGD2, and HLA-DPB1 in high glucose-treated THP-1 cells. qPCR results showed that the mRNAexpression levels of CXCL16, CTLA4, and HLA-DPB1 were significantly upregulated, while those of TGFBR1, CD48, and TMIGD2 were significantly downregulated in the high-glucose group compared to the normal glucose group (p < 0.05). TRIB1 knockdown significantly inhibited the abnormal expression of CXCL16, TGFBR1, CTLA4, CD48, TMIGD2, and HLA-DPB1 induced by high glucose (p < 0.05) (Fig. 10D). These results confirm the regulatory role of TRIB1 on immune-related genes in T1D.

The PI3K-AKT-mTOR signaling pathway is an important immune activation mechanism that has been shown to play a crucial role in the development of T1D in previous studies. In our bioinformatics analyses, marker genes of monocytes were significantly enriched in the PI3K-AKT signaling pathway, and the expression of key genes TRIB1, ACTG1, and REL was significantly correlated with the PI3K-AKT-mTOR signaling pathway. Therefore, we further investigated the effect of TRIB1 knockdown on the PI3K/AKT/mTOR signaling pathway in high glucose-treated THP-1 cells. Western blot results showed that there was no significant difference in the protein expression levels of PI3K, AKT, and mTOR between the high-glucose and normal glucose groups. However, the expression levels of p-PI3K, p-AKT, and p-mTOR were significantly upregulated in the high-glucose group. Compared with the high-glucose treatment group, TRIB1 knockdown significantly inhibited the phosphorylation of PI3K, AKT, and mTOR (p < 0.05) (Fig. 10E). These findings suggest that TRIB1 regulates the PI3K/AKT/mTOR signaling pathway in high glucose-treated monocytes, which may contribute to the immune dysregulation in T1D.

## 4 Discussion

T1D is a complex autoimmune disorder characterized by the destruction of insulin-producing pancreatic β cells, leading to hyperglycemia and severe complications ^[1]^. Despite advances in understanding the pathogenesis of T1D, the precise immune mechanisms underlying the disease remain elusive. In this study, we integrated scRNA-seq, bulk RNA-seq, and experimental validation to elucidate the immune mechanisms of T1D and identify potential diagnostic and therapeutic targets.

Our study identified monocytes as the key cell subtype with the most interactions with other cell subtypes in the immune landscape of T1D. This finding is consistent with previous studies that have highlighted the crucial role of monocytes in the pathogenesis of autoimmune diseases, including T1D ^[15, 16]^. Monocytes are innate immune cells that can differentiate into macrophages and dendritic cells, which play essential roles in antigen presentation, cytokine production, and the regulation of adaptive immune responses ^[17]^. In T1D, monocytes and macrophages have been shown to infiltrate the pancreatic islets and contribute to the inflammatory microenvironment that promotes β cell damage and apoptosis ^[15, 18]^. Monocytes and macrophages secrete proinflammatory cytokines, such as IL-1β and TNF-α, which can directly impair β cell function and survival ^[15, 19]^. Moreover, these cells can present autoantigens to T cells, leading to the activation and expansion of autoreactive T cells that mediate the destruction of β cells ^[15]^. Our study further emphasizes the central role of monocytes in orchestrating the immune responses in T1D and their potential as therapeutic targets. Modulating the activation and function of monocytes and macrophages may help attenuate the inflammatory response and preserve β cell mass in T1D.

Through the integration of scRNA-seq and bulk RNA-seq data, we identified 11 characteristic genes and constructed a diagnostic model with high validation efficiency. Among these genes, ACTG1, REL, and TRIB1 emerged as key players in T1D pathogenesis, showing consistent expression trends in the training and validation sets. ACTG1 encodes for gamma-actin, a cytoskeletal protein involved in cell motility and intracellular transport ^[20]^. While the specific role of ACTG1 in T1D has not been previously reported, cytoskeletal rearrangements and altered cell motility have been implicated in the activation and migration of immune cells in autoimmune diseases ^[20, 21]^. In T cells, cytoskeletal remodeling is crucial for immunological synapse formation and T cell receptor (TCR) signaling initiation. Dysregulation of cytoskeletal dynamics may contribute to abnormal T cell activation and function in T1D ^[22]^. REL, an NF-κB family transcription factor, is a well-established regulator of immune responses and has been linked to various autoimmune disorders, such as rheumatoid arthritis and multiple sclerosis ^[23]^. NF-κB signaling plays a critical role in the activation of immune cells and the production of proinflammatory cytokines. In T1D, increased NF-κB activity has been observed in immune cells and pancreatic islets, contributing to the inflammatory response and β cell apoptosis ^[24]^. Our study suggests that REL may also play a crucial role in the immune dysregulation observed in T1D, and targeting REL or NF-κB signaling may have therapeutic potential in modulating the inflammatory response in T1D. TRIB1, a member of the tribbles pseudokinase family, has been implicated in regulating cell proliferation, differentiation, and apoptosis ^[25]^. Recent studies have demonstrated TRIB1’s involvement in modulating immune responses and the development of autoimmune diseases, such as rheumatoid arthritis and inflammatory bowel disease ^[25–27]^. TRIB1 has been shown to regulate the differentiation and function of macrophages and T cells, two key cell types involved in T1D pathogenesis ^[28, 29]^. Our results highlight the potential role of TRIB1 in T1D pathogenesis and its regulation of immune-related pathways, suggesting that targeting TRIB1 may provide a novel therapeutic approach to modulate immune responses in T1D.

Functional analyses revealed that the key genes identified in our study were involved in immune regulation and the PI3K/AKT/mTOR signaling pathway. This pathway plays a pivotal role in various cellular processes, such as growth, proliferation, and metabolism, and has been implicated in the pathogenesis of several autoimmune diseases, including T1D ^[30, 31]^. In the context of T1D, the PI3K/AKT pathway has been shown to influence the activation and function of immune cells, contributing to the inflammatory response and β cell destruction ^[31–33]^. Activation of the PI3K/AKT/mTOR pathway in T cells can enhance their proliferation, survival, and effector functions, leading to exacerbated autoimmune responses ^[34]^. Meanwhile, in macrophages, this signaling pathway can modulate the production of proinflammatory cytokines and promote polarization towards an M1 phenotype, which is associated with increased inflammation and tissue damage ^[35]^. Interestingly, pharmacological inhibition of the PI3K/AKT/mTOR pathway has been demonstrated to mitigate the inflammatory response and improve disease outcomes in T1D animal models ^[31, 32, 36]^. Our findings further support the involvement of the PI3K/AKT/mTOR pathway in the immune dysregulation observed in T1D and highlight the potential therapeutic value of targeting this pathway. Modulating the activity of the PI3K/AKT/mTOR pathway in immune cells may help restore immune balance and prevent β cell destruction in T1D.

To validate our bioinformatics findings, we performed in vitro experiments using a high-glucose model in monocytes. High glucose induced monocyte activation, increased apoptosis, and altered the expression of key genes and immune-related genes, suggesting that hyperglycemia may contribute to the immune dysregulation in T1D. These findings are in line with previous studies that have demonstrated the deleterious effects of high glucose on immune cell function and the induction of inflammatory responses ^[37]^. Hyperglycemia has been shown to activate monocytes and macrophages, leading to increased production of proinflammatory cytokines and reactive oxygen species (ROS), which can contribute to β cell damage and the development of T1D complications ^[38, 39]^. Moreover, high glucose can induce epigenetic changes in immune cells, such as DNA methylation and histone modifications, which can alter gene expression and promote a proinflammatory phenotype ^[40, 41]^. Our study highlights the importance of glycemic control in modulating the immune responses in T1D and suggests that targeting the deleterious effects of hyperglycemia on immune cells may have therapeutic potential.

Notably, TRIB1 knockdown mitigated monocyte activation induced by high glucose conditions and influenced the expression of immune-related genes and the PI3K/AKT/mTOR pathway. These findings underscore the importance of TRIB1 in T1D pathogenesis and propose that TRIB1 could be a potential therapeutic target for modulating immune responses and reducing the harmful impact of hyperglycemia in T1D. Previous studies have demonstrated TRIB1’s role in regulating macrophage and T cell activation and function through various signaling pathways, such as MAPK and NF-κB ^[25, 42]^. In macrophages, TRIB1 promotes polarization towards an anti-inflammatory M2 phenotype and suppresses proinflammatory cytokine production ^[43]^. Furthermore, TRIB1 modulates CD4+ T cell activation and differentiation, as well as the Th1/Th2 balance in T cells ^[29]^. Our study expands on these findings, suggesting that TRIB1 may also play a crucial role in regulating monocyte activation and function in the context of T1D and hyperglycemia. As such, targeting TRIB1 could provide a novel therapeutic strategy for modulating immune responses and mitigating inflammation in T1D.

Although this research advances our understanding of T1D immunopathogenesis and identifies candidate molecules for diagnosis and treatment, several constraints should be acknowledged. The primary limitation relates to the restricted scale of transcriptomic analyses, necessitating validation in expanded patient populations. Additionally, the experimental framework, which utilized a monocytic cell line, may not fully represent the complexity of immune responses in primary human cells and animal models of T1D. Furthermore, the molecular mechanisms through which ACTG1 and REL influence T1D progression remain to be elucidated and require detailed examination. Notwithstanding these methodological constraints, this investigation establishes a robust framework for subsequent studies exploring T1D immunological mechanisms and the development of precision-based diagnostic tools and therapeutic interventions. Further research addressing these limitations will enhance our understanding of T1D pathogenesis and potentially lead to more effective treatment strategies.

In conclusion, our study integrates scRNA-seq, bulk RNA-seq, and experimental validation to unravel the immune mechanisms of T1D. We identified monocytes as the key cell subtype and ACTG1, REL, and TRIB1 as crucial genes involved in T1D pathogenesis. The constructed diagnostic model showed high validation efficiency and may have potential clinical applications. Laboratory investigations confirmed the role of TRIB1 in regulating monocyte activation and immune-related pathways in a high-glucose model, suggesting that targeting TRIB1 may have therapeutic potential in T1D. These discoveries enhance our comprehension of T1D’s immunological complexity and establish foundations for subsequent investigations and therapeutic innovations. By unraveling the intricate immune mechanisms underlying T1D, we hope to improve the diagnosis, prevention, and treatment of this debilitating autoimmune disorder. Future research should focus on developing targeted therapies that modulate the immune responses in T1D, such as small molecules or biological agents that inhibit the activity of key genes or pathways identified in our study. Furthermore, implementing comprehensive multi-omics approaches, encompassing transcriptomic, proteomic, and metabolomic analyses, could reveal deeper insights into the interactions between immune cells, β cells, and their microenvironment. Longitudinal studies that follow individuals at high risk for T1D may help elucidate the early immune events that precede the onset of clinical disease and identify novel biomarkers for early diagnosis and intervention. The ultimate objective is to translate these molecular insights into improved patient outcomes through enhanced understanding of T1D immunopathogenesis and the development of targeted therapeutic strategies.

## 5 Conclusions

In this study, we integrated scRNA-seq, bulk RNA-seq, and experimental validation to elucidate the immune mechanisms underlying type 1 diabetes (T1D). Through bioinformatics analyses, we identified monocytes as the key cell subtype with the most complex interactions with other immune cell subtypes in T1D. We also identified 11 characteristic genes and constructed a diagnostic model that demonstrated high validation efficiency in both training and testing cohorts. Notably, ACTG1, REL, and TRIB1 displayed consistent differential expression patterns between T1D patients and healthy controls, emerging as critical molecular players in disease development. Pathway analysis demonstrated that these essential genes participated in immune system regulation and PI3K/AKT/mTOR pathway signaling. In vitro experiments using a high-glucose model in the monocytic cell line THP-1 confirmed that high glucose induced monocyte activation, as evidenced by increased expression of activation markers (CD86) and pro-inflammatory cytokines (IL-8 and TNF-α). High glucose also increased monocyte apoptosis and altered the expression of key genes (ACTG1, REL, and TRIB1) and immune-related genes (CXCL16, TGFBR1, CTLA4, CD48, TMIGD2, and HLA-DPB1). Significantly, TRIB1 suppression via siRNA reduced glucose-mediated monocyte activation and regulated immune-related gene expression and PI3K/AKT/mTOR signaling, indicating its significance in modulating monocyte function and immune responses in T1D.

Our findings provide novel insights into the immune mechanisms of T1D, highlighting the importance of monocytes and the key genes ACTG1, REL, and TRIB1 in disease pathogenesis. The constructed diagnostic model based on the characteristic genes shows potential for clinical application in T1D diagnosis. Moreover, the recognition of TRIB1 as a key modulator of monocyte activation and immune pathways under hyperglycemic conditions suggests its potential as a therapeutic candidate. Additional research is warranted to confirm these observations in expanded patient populations and evaluate the therapeutic implications of targeting TRIB1 and related molecules in T1D treatment.

